# Wheat fiber-induced peripheral regulatory T-cells suppress development of colitis

**DOI:** 10.1101/2025.03.31.646427

**Authors:** Seong-eun Kim, Hirohito Abo, Yanling Wang, Shawn Winer, Dan Winer, Michael Pellizzon, Vu L. Ngo, Andrew T. Gewirtz

## Abstract

Reduced dietary fiber intake is associated with, and may have contributed to, the post-mid-20^th^ century increase in immune-mediated chronic inflammatory diseases, including inflammatory bowel disease. Reduced fiber intake has resulted, in part, from highly refined foods. For example, modern methods of producing bread removes much of the fiber naturally present in wheat kernels. Accordingly, we hypothesized that wheat fiber might protect against chronic inflammatory diseases. We tested this notion in a murine T-cell transfer colitis model. *Rag1^-/-^* mice were fed open-source low-fiber diets enriched, or not with wheat fiber (WF) and then administered CD45Rb^hi^ T-cells. WF conferred robust protection in this colitis model as assessed by an array of clinical, histopathologic, morphologic, and immune-related parameters. WF’s protection against colitis associated with a microbiota-dependent increase in Foxp3^+^ T-cell (Tregs), which could be recapitulated in vitro. WF did not induce Tregs in *CNS1^-/-^* mice nor did WF protect against T cell transfer colitis driven by transplant of colitogenic T-cells from *CNS1^-/-^* mice. Thus, enriching diet with WF has potential to promote microbiota-dependent peripheral Treg development and, consequently, protect against chronic inflammatory diseases.

## Introduction

Intertwined changes in food production and dietary habits have resulted in decreased consumption of dietary fiber in a manner that roughly parallels the increasing prevalence of immune-mediated chronic inflammatory diseases like inflammatory bowel disease (IBD)^1–3^. This correlation, combined with the fact that dietary fiber directly interacts with, and is metabolized by, gut microbiota, which is increasingly recognized to be a key chronic inflammatory disease determinant, has led to the hypothesis that fiber consumption protects against an array of diseases including IBD. In accord with this notion, maintaining mice on low-fiber diets, rather than the standard fiber-rich grain-based chow diets that lab mice are typically fed, increases severity of colitis in both acute (DSS) and T-cell transfer colitis models. Subsequent attempts to ameliorate colitis severity by enrichment of low-fiber diets with specific semi-purified fibers found fiber- and model-specific impacts. For example, inulin, which is readily fermentable to butyrate, which is known to induce regulatory T-cells (Treg), provided modest protection in a T-cell mediated colitis model^4^ but markedly exacerbated DSS-induced colitis^5^. In contrast, psyllium, which is only minimally fermentable, conferred strong protection in both the DSS and T-cell transfer model^6^. Psyllium’s protection was mediated activation of the FXR bile acid receptor, while inulin’s exacerbation of DSS colitis occurred despite it increasing bile acid levels and activating this receptor.

The rationale for our initial focus on these particular fibers is that many humans seek to improve their health by consuming them in supplements. However, their dramatically differing impacts on microbiota and colitis severity led us to appreciate the importance of also examining impacts of fibers naturally present in foods, the reduced consumption of which correlates with increased chronic inflammatory disease prevalence. Hence, this study, and a sister study performed in parallel, examined impacts of wheat fiber (WF). Wheat kernels, particularly the bran portion, are rich in fiber such that “whole wheat” or “whole-grain” breads can approach 15% fiber by weight. The majority of this fiber is absent in “white” breads made from the endosperm portion of the wheat kernel. Thus, we compared the consequence, in mice of maintaining them on low-fiber diets enriched, or not, with WF. Our parallel study found that WF protected against DSS colitis via an FXR-independent microbiota-dependent macrophage-mediated mechanism. Here, we report that WF protected against T-cell transfer colitis via microbiota-mediated generation of non-SCFA metabolites that promoted peripheral Treg development.

## Results

### Wheat fiber protects against T cell-mediated colitis

To probe if wheat fiber (WF), which protects mice against acute colitis, might also ameliorate T-cell-mediated colitis *Rag1^-/-^* mice, raised on grain-based chow (GBC) were fed a low-fiber (LF) compositionally defined or WF-enriched version of this diet (Table 1). Two weeks later, such mice were administered CD4^+^CD25^-^CD45Rb^hi^ T (i.e. naïve) cells, freshly isolated from GBC-fed WT mice (Fig. 1A, Fig. S1). Body weight and fecal lipocalin-2 (Lcn2) levels, which reflect intestinal inflammation^7^, were monitored. LF-fed mice exhibited indices of colitis, including weight loss, elevated fecal Lcn2 expression by 5 weeks post-T cell transfer (Fig. 1B-C) and, a few weeks later, increased disease activity index (Fig. 1D) while approaching, or in the case of one mouse reaching, endpoints a few weeks later while WF-fed exhibited only modest changes in these parameters suggesting WF protected against colitis in this model. Post-euthanasia analysis confirmed this notion. Specifically, compared to LF-fed mice, WF-fed mice exhibited longer colons (Fig. 1E), smaller spleens (Fig. 1F). reduced TNFα expression (Fig. G), and lower histopathologic disease scores (Fig. 1H). Enrichment of diet with WF also impacted developmental fate of the transplanted CD4 T cells. Specifically, WF-fed mice displayed lower levels of Ifnɣ^+^, Il17a^+^, and Tnfα^+^ CD4 T cells and increased levels of FoxP3^+^ regulatory T cells (Tregs) (Fig. 1I), all of which might have contributed to the reduced disease exhibited by WF-fed mice.

**Fig. 1.**
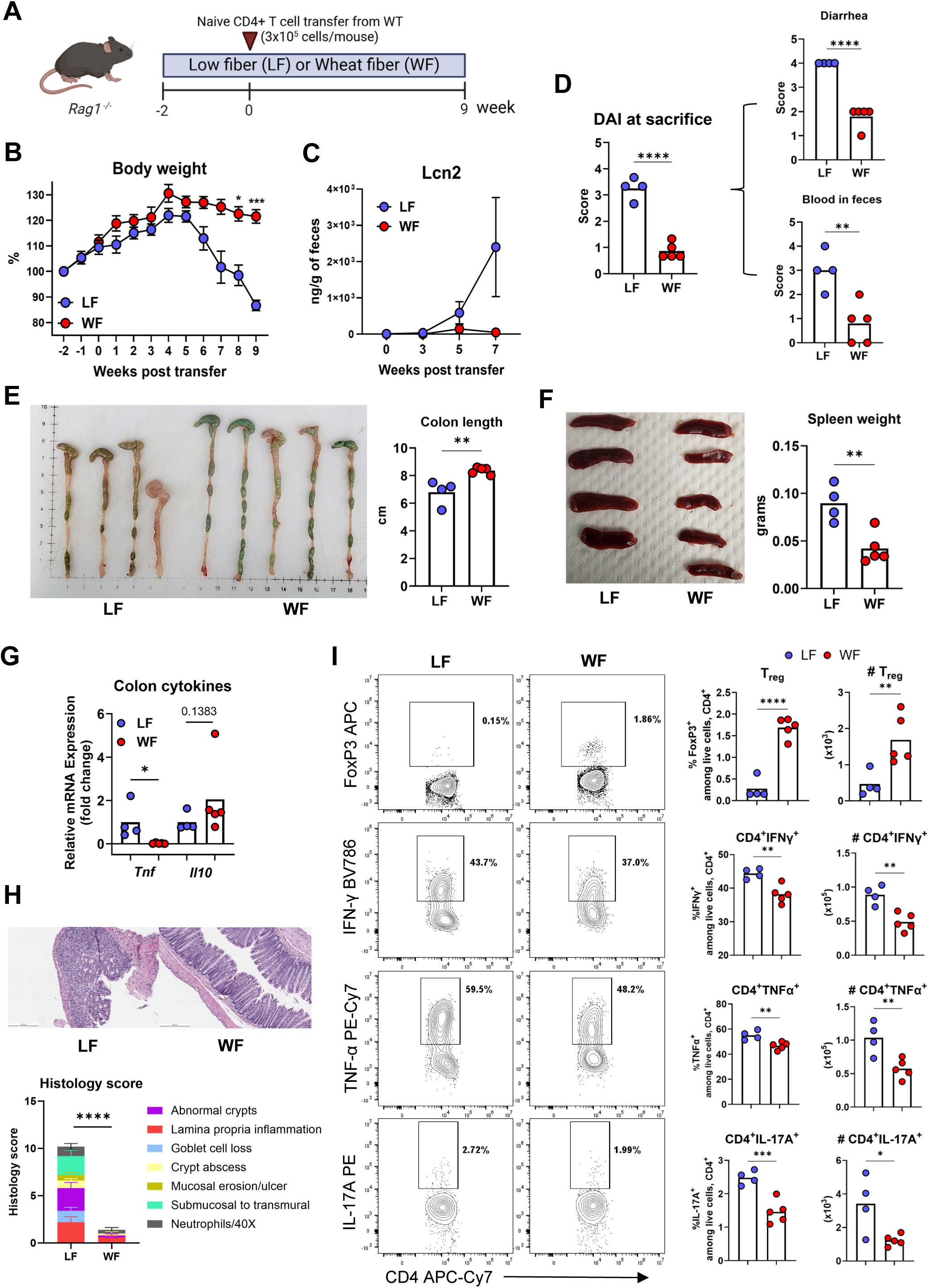
Wheat fiber protected mice against T-cell transfer colitis. Female *Rag1^-/-^* mice (n=5 per group) were fed either a low-fiber or wheat fiber-enriched (Wheat) diet for 2 weeks and received 3x10^5^ naïve CD4^+^ T cells from WT mice. (A) Schematic. (B) Body weight. (C) Fecal Lcn2 levels measured by ELISA. (D) Disease activity index. (E) Colon length and spleen weight at sacrifice. (G) Colon pro-inflammatory cytokines assessed by RT-qPCR. (H) Representative colon histology and scoring. (I) Colon lamina propria CD4^+^ T cells analyzed by flow cytometry.

**Fig. 2.**
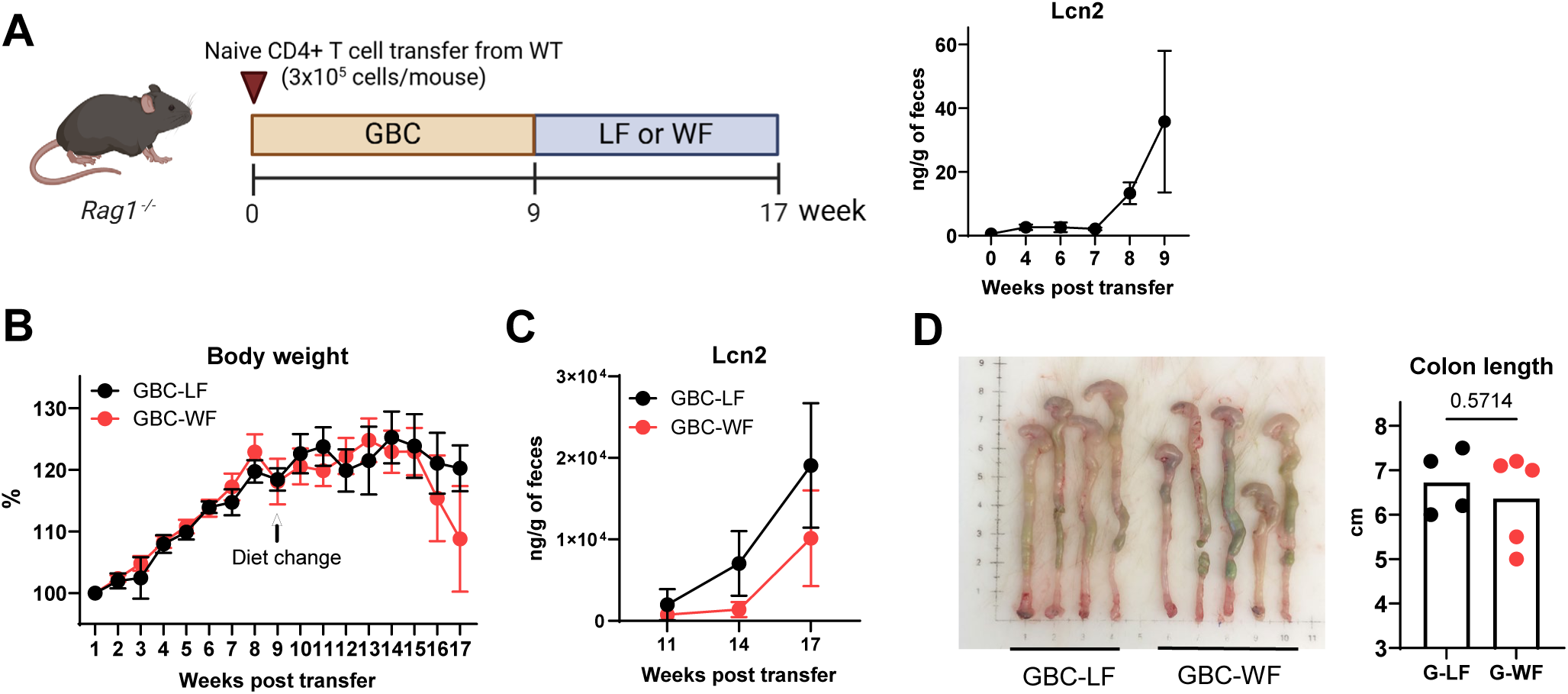
Wheat fiber did not ameliorate T-cell colitis following disease onset. *Rag1^-/-^* mice (n=4-5 per group) were injected with naïve CD4^+^ T cells while on GBC diet, then switched to either a low-fiber (LF) or wheat fiber (Wheat) diet. (A) Experimental design schematic. (B) Body weight and fecal Lcn2 levels post-transfer before diet change. (C) Fecal Lcn2 levels after diet change (D) Colon length at sacrifice.

**Fig. 3.**
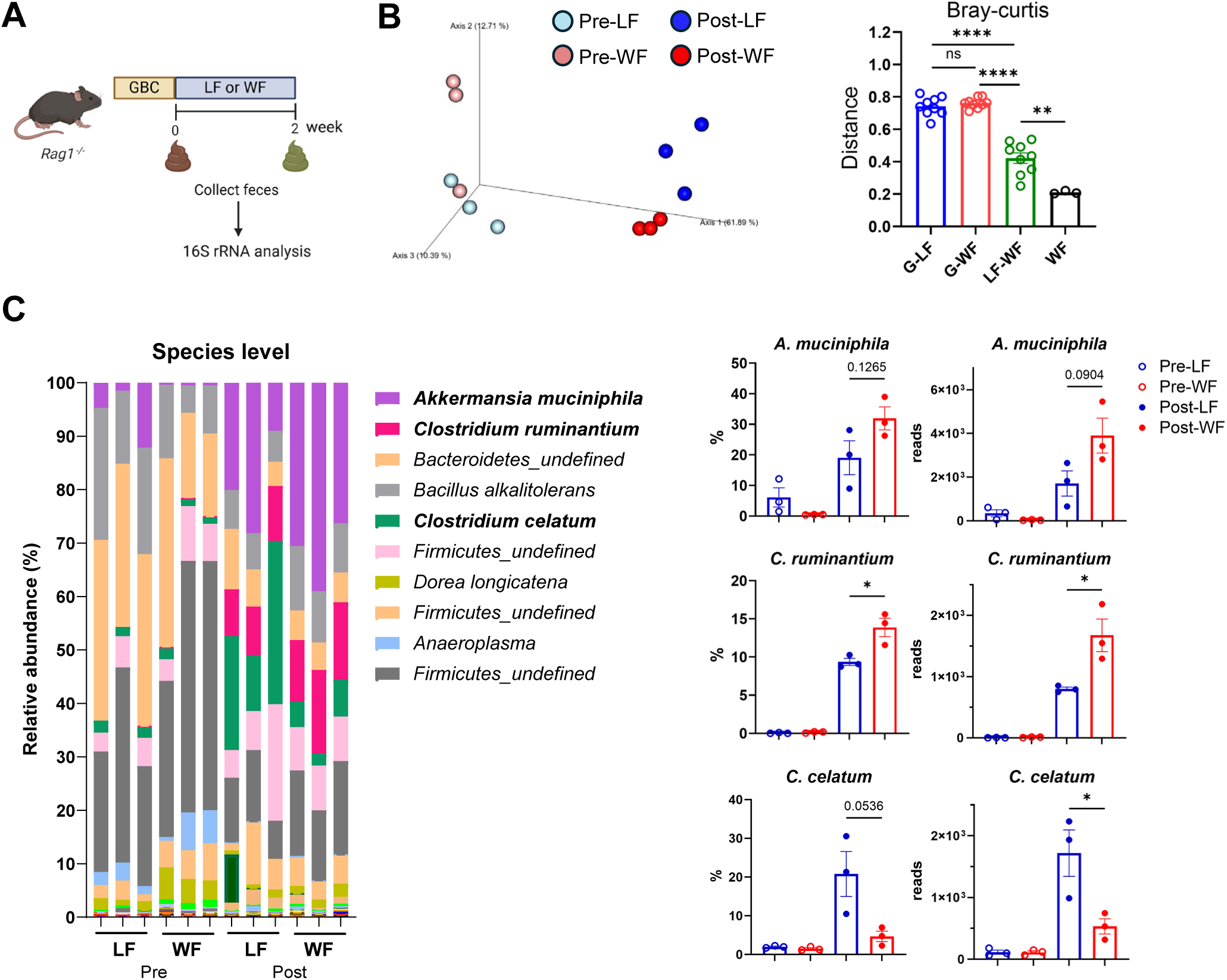
Wheat fiber altered gut microbiota composition in *Rag1^-/-^* mice. 16S rRNA sequencing was performed on feces of *Rag1^-/-^* mice fed either LF or Wheat for 2 weeks. (A) Schematic (B) Beta diversity (Jaccard) analysis (C) Taxonomic composition at the species level. (D) Three species altered by wheat fiber.

**Fig. 4.**
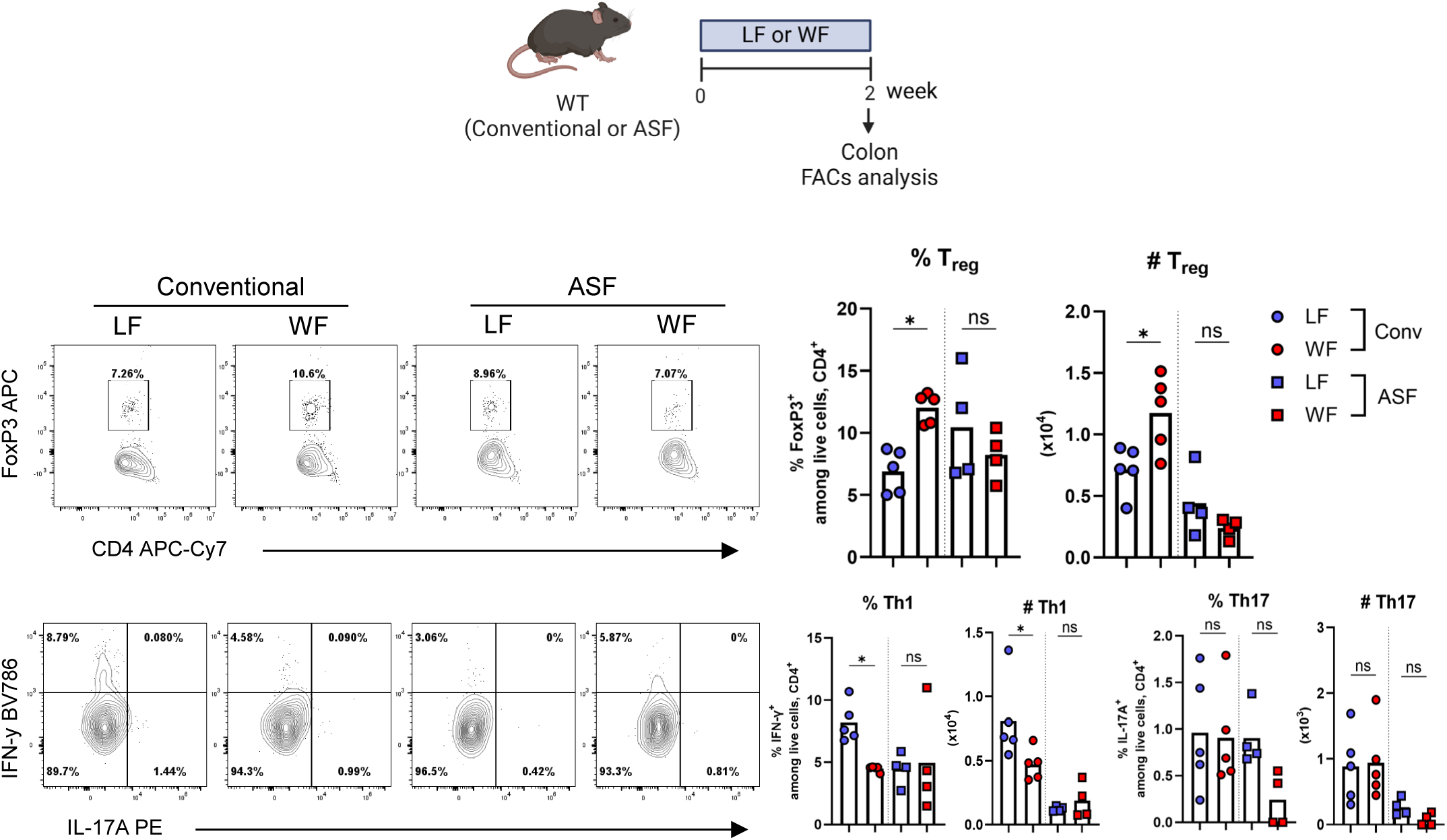
Wheat fiber increased regulatory T cells in a microbiota-dependent manner. Colonic lamina propria CD4^+^ T cells from different mice on indicated diets were measured by flow cytometry. (A) Regulatory T cells in conventional WT mice fed GBC or LF or Wheat for 2 weeks. (B) Regulatory T cells, Th1, and Th17 cells in conventional mice and ASF mice fed either LF or Wheat for 2 weeks.

We also examined the extent to which WF might impact colitis in mice with established, but early-stage, T-cell mediated colitis. GBC-fed *Rag1^-/-^* mice were administered CD4^+^CD25^-^ CD45Rb^hi^ T cells. Monitoring them by body weight and fecal Lcn2 levels suggested colitis had developed by 9-weeks post T-cell transplant, at which time mice were switched to the LF or WF diet. Body weight stabilized in both groups potentially reflecting that the dietary change resulted in a rapid change in microbiota-derived T-cell antigens but resumed a few weeks later at which point WF-fed mice developed colitis at least as severe as LF-fed mice as evidenced by all parameters assayed. Collectively, these results accord with the possibility that WF consumption might prevent but is unlikely to help manage T-cell mediated colitis.

### Wheat fiber increases regulatory T cells in a microbiota-dependent manner

Many impacts of dietary fiber, including WF’s protection against DSS colitis, are associated with, and require, alterations in gut microbiota, prompting us to examine how dietary changes impacted the microbiotas of *Rag1^-/-^* mice prior to induction of T-cell mediated colitis. Analogous to studies in WT mice, switching diet of *Rag1^-/-^* mice from GBC to either the LF or WF resulted in a stark change in microbiota composition. Differences in microbiota composition between LF and WF fed mice were more moderate but nonetheless PCoA analysis indicated a significant difference in overall composition and some specific taxa including WF enrichment of *Clostridium ruminantium* and depletion of *Clostridium celatum*, as well as a trend toward increased *Akkermansia muciniphilia* without impacting overall α-diversity (Fig. S2B). The absolute dependence of the CD45Rbhi T-cell transfer colitis model on gut microbiota precludes the use of gnotobiotic mice to examine if WF’s protection in this model required microbiota. Thus, to gain insight into this question, we examined if WF might impact T-cell differentiation, which is known to be a major determinant of disease severity in T-cell transfer colitis models. Specifically, we analyzed colonic CD4^+^ T-cells in mice following 2 weeks feeding with LF or WF diet. WF resulted in a significant increase in the percentage and absolute number of Tregs and a reduction in Th1 cells while Th17 cells remained unchanged. The level of Tregs in WF-fed mice was similar to that of inulin-fed mice and exceeded those of GBC-fed mice (Fig. S3). In contrast to inulin, which increases Tregs via SCFA generation^8^, the WF-induced increase in Treg was not associated with increased butyrate production (DSS ref) arguing against a role for fermentation. To assess whether WF-induced such changes in T-cell required microbiota, we turned to Altered Schaedler Flora (ASF) mice, which harbor a minimal eight species microbiota^9^ that, unlike germ-free conditions, confer relatively normal immune function. WF feeding did not impact Tregs or Th1 cells in ASF mice. Thus, WF promotion of Treg development is microbiota-mediated.

### WF-induced Tregs mediate its protection against T-cell mediated colitis

The results outlined above led us to hypothesize that feeding WF-enriched diets to *Rag1^-/-^* resulted in an environment that favored transplanted naïve T-cells becoming Tregs. In thinking about such an environment, we first hypothesized a role for M2-like macrophages, which are known to promote Tregs and are increased by WF in WT mice (cite DSS paper). However, we did not observe increased colonic levels of F4/80^+^CD206^+^ cells (i.e. M2-like macrophages) in response to WF in *Rag1^-/-^* mice (Fig. 5A) arguing against this possibility. Hence, we next considered the possibility that WF feeding might generate colonic metabolites that directly acted on naïve CD4 T-cells to favor Treg development. We investigated this possibility via a well-established in vitro assay wherein naïve CD4^+^ T cells are incubated in anti-TCR coated plates in cytokines that enable differentiation into Tregs. We added the fecal supernatants (FS) of *Rag1^-/-^* mice fed LF or WF to this environment. While FS from mice fed either diet impeded Treg development in this assay, the extent to which it did so was less in WF FS, resulting in Tregs being more abundant in WF vs. LF conditions (Fig. 5B) suggesting that a WF-derived component of FS favors Treg development. The *in vitro* differentiation assay we utilized is thought to yield cells reminiscent of peripheral Tregs, whose development is dependent upon conserved non-coding sequence 1 knock-out (*CNS1*)^10^. Accordingly, naïve CD4 T-cells from *CNS1^-/-^* mice did not develop into Tregs in vitro, even in the absence of any inhibitory activity present in FS. Moreover, WF feeding to *CNS1^-/-^* mice did not increase levels of colonic Tregs supporting our presumption that WF was promoting Treg development in the periphery.

**Fig. 5.**
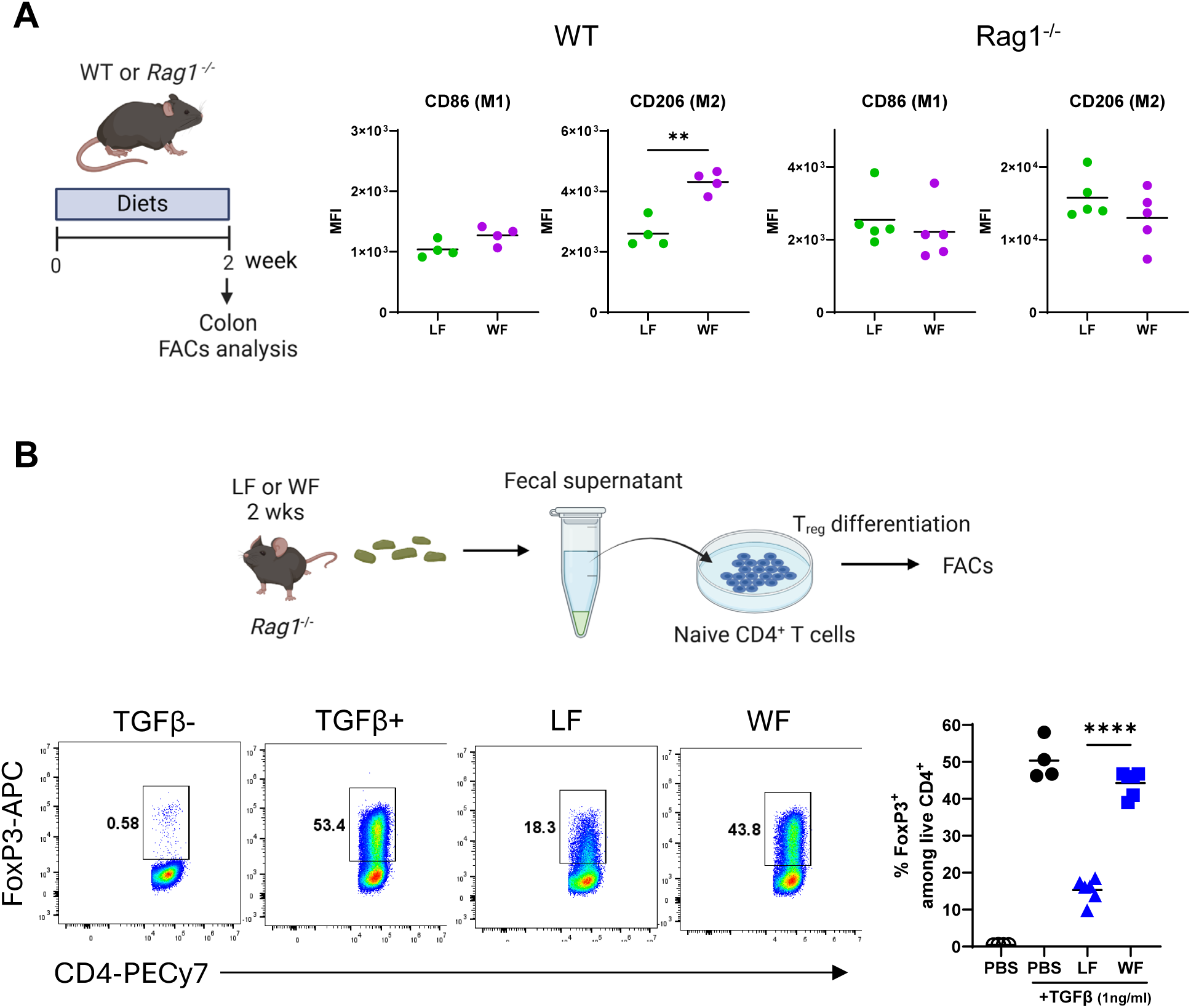
Wheat fiber didn’t change the innate immune cells in *Rag1^-/-^* mice but fecal supernatant from wheat fiber-fed mice can induce regulatory T cells *in vitro*. (A) WT or *Rag1^-/-^* mice (n=3-5 per group) were fed with GBC or LF or Wheat for 2 weeks and their colon macrophages were analyzed by flow cytometry. (B) Naïve CD4^+^ T cells from WT mice were cultured for 3 days on aCD3ε-coated plates (0.75– 1x10⁵ cells/well) with aCD28 (2.5 µg/ml), IL-2 (15 U/ml), and TGF-β (2 ng/ml), except for the negative control, which lacked TGF-β. Fecal supernatants from LF or Wheat-fed *Rag1^-/-^* mice were added. Flow cytometry analysis of induced Tregs with fecal supernatants.

**Fig. 6.**
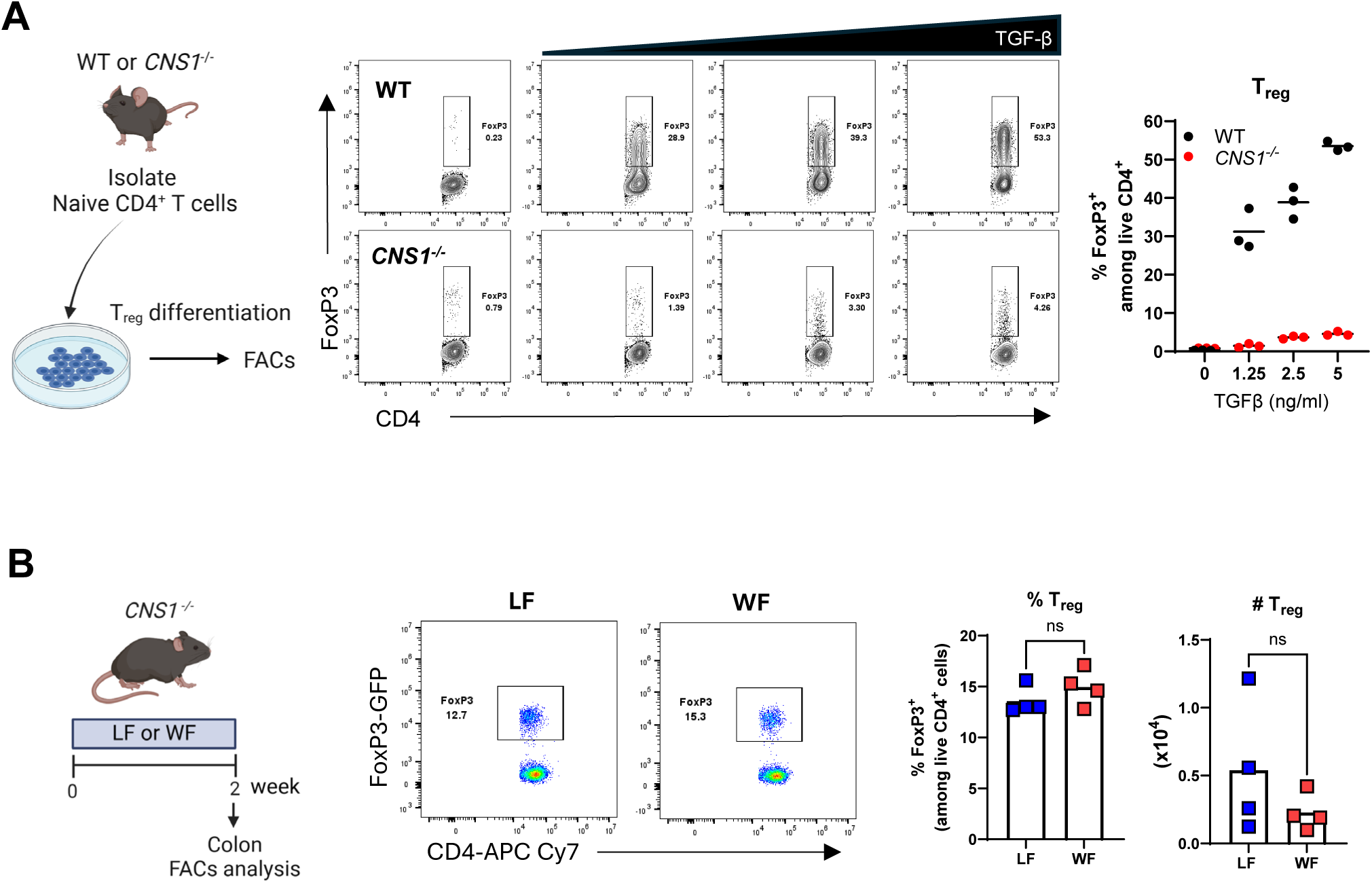
Wheat fiber did not increase regulatory T cells in *CNS1^-/-^* mice, whose naïve T cells have impaired induction of Tregs. (A) Naïve CD4^+^ T cells were isolated from WT or *Rag1^-/-^* mice and cultured in vitro with various concentrations of TGF-β for 3 days and Tregs were analyzed by flow cytometry. (B) *CNS1^-/-^* mice were fed with either LF or Wheat for 2 weeks and their colonic lamina propria Tregs were analyzed by flow cytometry.

**Fig. 7.**
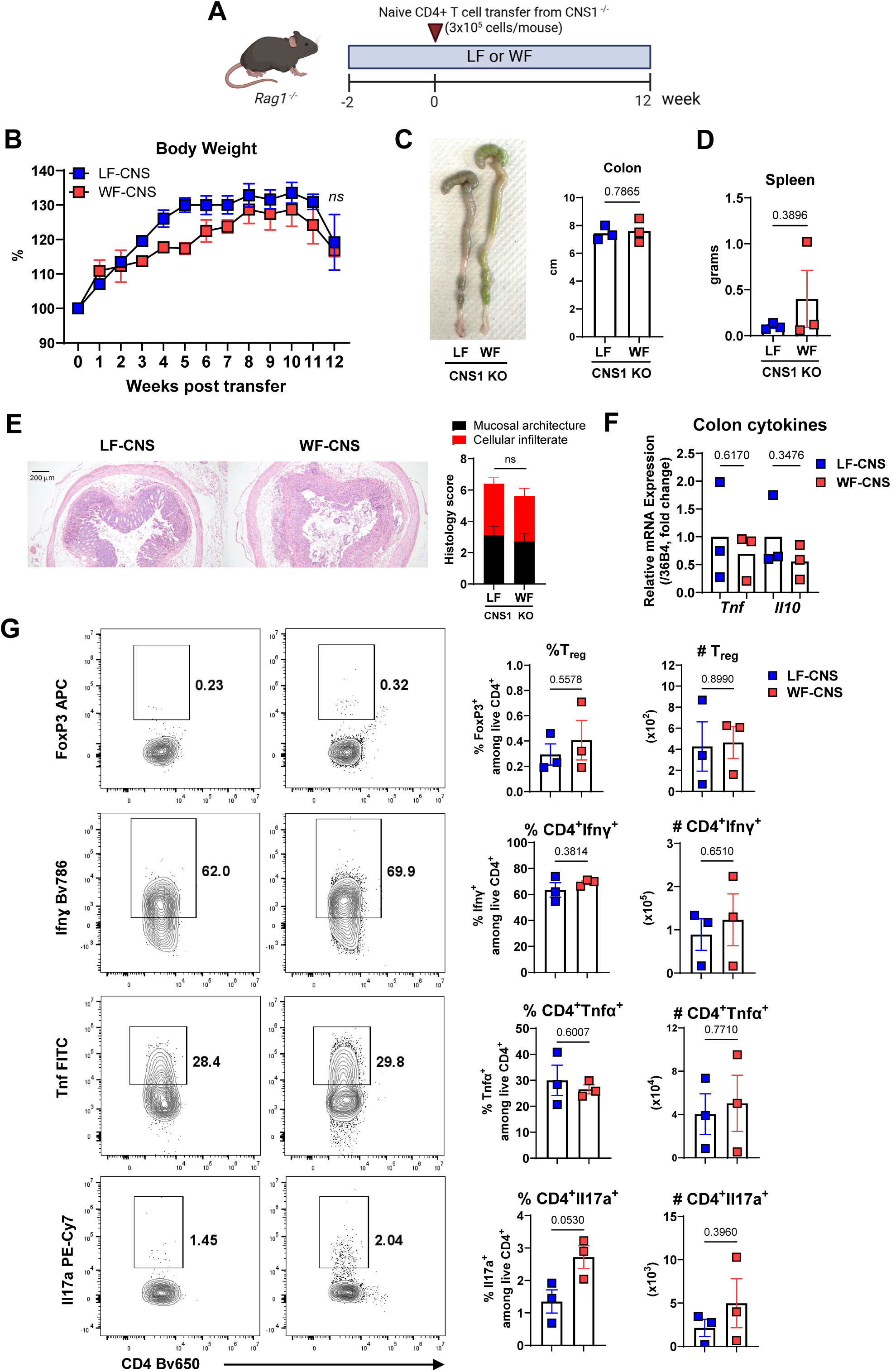
Wheat fiber did not protect against T cell colitis in the absence of peripheral regulatory T cells. *Rag1^-/-^* mice (n=5 per group) were fed either LF or Wheat for 2 weeks and received splenic 3x10^5^ naïve CD4^+^ T cells from *CNS1^-/-^* mice. (A) Body weight. (B) Spleen weight. (D) Colon length. (E) Colon inflammatory cytokine levels measured by RT-qPCR. (F) Representative colon histology and scoring. (G) Colonic lamina propria CD4^+^ T cells analyzed by flow cytometry.

Lastly, we probed the role of peripheral Treg development in mediating WF’s protection against T-cell mediated colitis. CD4^+^CD25^-^CD45Rb^hi^ (i.e. naïve) T-cells, isolated from GBC-fed *CNS1^-/-^* mice, were transplanted into *Rag1^-/-^* mice consuming LF or WF diets. Both groups of mice began to lose weight 10 weeks post-transfer and did so with similar kinetics suggesting a lack of protection by WF. Moreover, post-mortem assessment of colitis, based on morphology, histopathology, and cytokine analysis indicated WF did not protect against colitis driven by *CNS1^-/-^* T-cells. Furthermore, WF feeding did not result in increased levels of FoxP3^+^ cells, nor did it decrease levels of Ifnɣ^+^, Tnfα^+^, or Il17a^+^ CD4 T cells. Thus, the ability of WF feeding to promote Tregs, suppress colitogenic T-cells, and protect mice against T-cell mediated colitis required *CNS1*.

## Discussion

The dearth of fiber in modern diets is correlated with increased prevalence of chronic inflammatory diseases, including IBD^1–3^. Although the extent to which this reflects causation is not yet clear. Recent studies suggest that the extent to which specific dietary fibers impact colitis is highly context-dependent^11^. In this study, we demonstrated that wheat fiber, historically a staple but largely diminished following dietary modernization, plays a crucial role in maintaining gut immune homeostasis and preventing mice from severe colitis by promoting development of Treg. Wheat fiber provided strong protection against colitis, when T cell-deficient mice were fed wheat fiber prior to the transfer of naïve T cells. However, this protective effect was lost when *CNS1^-/-^* naïve T cells, incapable of differentiating into Tregs, were transferred. Given the essential role of Tregs in suppressing inflammation and autoimmune responses, our findings suggest that the decline in dietary fiber intake may have contributed to increased disease susceptibility. Previous research has shown that wheat-derived arabinoxylan, can enhance butyrate production and Treg differentiation, thereby conferring protection in this adoptive transfer model of colitis^12^. The specific wheat fiber used here was low in arabinoxylans and was not highly fenentable and thus likely drove Tregs by a distinct mechanism. Notably, wheat fiber supplementation after disease onset did not confer the same protective effect, indicating its benefits may depend on long-term dietary intake prior to severe inflammation. Further research is needed to determine the precise timing for wheat fiber to confer protection before and after disease onset. Additionally, these findings may imply that the benefits of naturally occurring dietary fiber lost over time may not be readily restored by simply increasing wheat fiber intake within a short period.

Wheat fiber increased peripheral Tregs in a microbiota-dependent manner. Most Tregs in the colon are pTregs as naive CD4 T cells are constantly exposed to different microbial stimuli. Given fecal supernatant from wheat fiber-fed mice can induce more Tregs, we also presume that some wheat fiber-induced metabolites may be responsible for promoting Tregs. We acknowledge that we do not know the exact component of wheat fiber that drives pTregs. SCFA such as butyrate, typically a result of fermentable fiber, are known to induce Tregs via GPR 43 or GPR109^13, 14^, but our wheat fiber is mostly (97%) insoluble, non-fermentable fiber, producing a similar amount of butyrate in mice cecum compared to the low-fiber diet suggesting other metabolites are likely to be involved. Most studies of dietary fiber study have focused on fermentable fibers and their metabolites as insoluble, non-fermentable fibers are thought to be merely helping motility and producing bulky feces. However, our result indicated that insoluble wheat fiber could still change the gut microbiota and Tregs in a microbiota-dependent manner.

Although wheat fiber didn’t change the diversity of gut microbiota, wheat fiber still brought remarked change in several species: *A. muciniphila* could be one of the candidates that’s responsible for wheat-fiber-induced pTregs, as the bacteria’s epitope was previously shown to induce Treg^15^. The abundance of this bacteria is low in IBD patients^16^ and it has been correlated with protective roles in IBD^17^, but further study is needed to define the keystone for pTreg-derived protection. Wheat fiber also increased *Clostridium ruminantium* which is typically found in ruminants^18^ and has not yet been reported in the murine gut. The abundance of *Clostridium celatum* was lowered by wheat fiber, and a similar trend was observed by a plant-derived anti-inflammatory flavonoid ^19, 20^. Also, this bacteria was increased in the groups with diarrhea in immunocompromised mice^21^ and celiac disease patients^22^.

Future research should focus on identifying the key wheat fiber-derived microbial metabolites responsible for pTreg differentiation, as well as determining the optimal timing and duration of fiber intake for long-term immune benefits. Additionally, clarifying the metabolites that drive Treg development may yield novel strategies to prevent this disorder.

## Methods

### Mice

C57BL/6 wild-type (WT) and *Rag1^-/-^* were purchased from Jackson Laboratories (Bar Harbor, ME) and Conserved Non-coding Sequence 1 knockout (*CNS1^-/-^*) mice were provided by Alexander Rudensky (Sloan Kettering Institute). Gnotobiotic mice with ASF were generated from C57BL/6 germ-free mice purchased from Taconic Biosciences Inc. (Rensselaer, NY). All mice were bred/housed at Georgia State University under Institutional Animal Care and Use Committee #A20043 and #A24001. Both male and female mice were used in the preliminary and found no difference thus female mice were used throughout the experiment. All mice were maintained under normal grain-based chow (GBC) until the indicated experimental diets: a low-fiber diet (LF), containing only 50 grams of cellulose as a dietary fiber or LF enriched with 150 grams of purified wheat fiber.

### Adoptive T cell transfer model of colitis

Nine-week-old *Rag1^-/-^* mice were administered FACS-sorted CD4^+^CD25^-^CD45Rb^hi^ T cells (2-3x10^5^ cells/mouse) via intraperitoneal injection. Such naïve T cells were isolated from spleens of either C57BL/6 WT mice or *CNS1^-/-^* mice as indicated. After the transfer, the recipient mice were monitored weekly for weight loss and diarrhea and euthanized when mice lost >20% of body weight on the day of transfer. The indicated diets were fed either before or after the transfer of naïve T cells.

### Isolation of colonic lamina propria (colon digestion)

Dissected colons were cut into small pieces and washed with calcium- and magnesium-free Hanks’ balanced salt solution (CMF/HBSS). The epithelium was removed by two consecutive incubation in a shaker (200 rpm, 20 min, 37 °C) with 2 mmol/L ethylenediaminetetraacetic acid (EDTA) in CMF/HBSS containing 5% fetal bovine serum. Intestinal tissue pieces were digested with 1 mg/ml of collagenase type IV (Sigma C5138) and 40 μg/ml of DNase I (Roche, Germany) in CMF/HBSS or RPMI containing 5% fetal bovine serum at 37 °C for 10-12 minutes. Cells were then filtered (40μm), centrifuged and supernatants were discarded and resuspended in cold CMF/HBSS. Lamina propria mononuclear cells were further purified by Percoll (GE Health Care, Uppsala, Sweden) density gradient centrifugation.

### In vitro differentiation of regulatory T cells

Naïve CD4⁺ T cells were isolated from the spleens of wild-type (WT) mice using the MACS purification kit (Miltenyi Biotec, #130-104-453) following the manufacturer’s instructions. The day before, a 96-well cell culture plate was coated overnight with anti-CD3 antibodies (2.5 µg/mL). Purified naïve CD4⁺ T cells (0.8∼1 × 10⁵ per well) were then cultured for three days in T cell medium (RPMI, 10% fetal bovine serum, 1% HEPES, 1% nonessential amino acids, 1mM sodium pyruvate, 55µM 2-mercaptoethanol, 100 U/ml penicillin, 100 µg/ml streptomycin) supplemented with anti-CD28 (1 µg/mL), recombinant human IL-2 (15 U/mL), and recombinant human TGF-β (1–5 ng/mL). Fecal supernatants were added as indicated in the corresponding figures.

### Flow cytometry

Cells were first blocked with 1 μg per million cells of 2.4G2 (BioXCell) to prevent Fc receptor-mediated binding. Live cells were identified by excluding dead cells using the Fixable Aqua Dead Cell Staining Kit (Invitrogen). After washing with PBS, cells were stained for surface markers (i.e., CD4) using conjugated monoclonal antibodies (mAbs) in FACS buffer. For intracellular staining, cells were re-stimulated with 1X Cell Activation Cocktail with Brefeldin A (BioLegend) for 3 hours, followed by fixation and permeabilization using the Fixation/Permeabilization (Fix/Perm) Buffer Set (eBioscience). Subsequently, cells were stained for inflammatory cytokines and FoxP3 in the FoxP3 Staining Buffer (eBioscience). Fluorochrome-conjugated mAbs were used as indicated in the figures. Multi-parameter flow cytometry was performed on a CytoFlex (Beckman Coulter) and analyzed using FlowJo software (Tree Star). Cell sorting was conducted on a SONY SH800Z cell sorter (Sony).

### RNA extraction and qRT-PCR

Total RNA was extracted from colons using Trizol (Invitrogen, Carlsbad, CA) according to the manufacturer’s protocol. Quantitative RT-PCR was performed using iTaq™ One-Step RT-PCR Kit with SYBR Green (Bio-Rad, Hercules, CA) in a CFX 96 apparatus (Bio-Rad, Hercules, CA). using the following primers (F/R): *36b4*: 5’- TCCAGGCTTTGGGCATCA-3’, 5’- CTTTATTCAGCTGCACATCACTCAGA-3’; *Tnf*: 5’-CGAGTGACAAGCCTGTAGCC-3’, 5’- CATGCCGTTGGCCAGGA-3’; *Il10*: 5’-CAGTACAGCCGGGAAGACAA-3’, 5’- GGCTTGGCAACCCAAGTAA-3’. Differences in transcript levels were normalized to the housekeeping gene *36b4*.

### Histopathologic analysis

Fresh colons were fixed in 10% PBS-buffered formalin. Paraffin embedding, sectioning, and hematoxylin and eosin (H&E) staining were performed at HistoWiz, Inc. The stained slides were blindly scored by two pathologists, Shawn Winer. and Daniel A. Winer.

### Bacteria quantification in feces by qPCR

To measure the total fecal bacterial load, total DNA was isolated from weighted feces using QIAamp DNA Stool MiniKit (Qiagen, Hilden, Germany). DNA was then subjected to qPCR using QuantiFast SYBR Green PCR kit (Bio-Rad, Hercules, CA) with universal 16S rRNA primers 8F: 5’-AGAGTTTGATCCTGGCTCAG-3’ and 338R: 5’- CTGCTGCCTCCCGTAGGAGT-3’ to measure total bacteria number, and bacteria-specific primers (*B.theta* F: 5’-TACTCGCCTCTTTGCAACCCTACC-3’, R: 5’- GGCCCCAGATCCGAACAACAC-3’ and *A. muciniphila* F: 5’- CAGCACGTGAAGGTGGGGAC-3’, R: 5’-CCTTGCGGTTGGCTTCAGAT-3’). Results are expressed as bacteria number per mg of stool using a standard curve.

### Lipocalin-2 ELISA

Fecal lipocalin-2 levels were measured using DuoSet mouse lipocalin-2 ELISA kit (R&D Systems, #DY1857) according to the protocol. Fecal supernatants were collected after centrifuge (12,000 rpm, 10 min, 4 °C) of the homogenized feces (100 mg of feces/ml of PBS).

### Gut microbiota analysis by 16S rRNA gene sequencing

Fecal DNA was extracted using DNeasy 96 PowerSoil Pro QIAcube HT Kit (Qiagen), and the region V3-V4 of 16S rRNA genes were amplified using the following primers: 341F 5′- TCGTCGGCAGCGTCAGATGTGTATAAGAGACAGCCTACGGGNGGCWGCAG-3′; 805R 5′- GTCTCGTGGGCTCGGAGATGTGTATAAGAGACAGGACTACHVGGGTATCTAATCC- 3′. PCR products of each sample were purified using Ampure XP magnetic purification beads. An index PCR was performed to attach dual barcodes and Illumina sequencing adapters using Nextera XT Index kit (Illumina). Final PCR products were verified on 1.5% DNA agarose gel and quantified using Pico dsDNA assay (Invitrogen). An equal molar of each sample was combined and purified again using Ampure XP beads as the library. The library was diluted and spiked with 5% PhiX control (Illumina) and sequenced by Illumina MiSeq sequencing system (2 × 250bp). Demultiplexed fastq files were generated on instrument. Sequence reads were quality-filtered by DADA2 plugin in Qiime2. Taxonomy was assigned based on the Greengenes database. Raw sequencing data has been deposited at the website(URL) with accession number: 000000.

### Statistical analysis

GraphPad Prism 10.0 was used for all statistical testing. All data are given as mean ±SEM, and were compared via Student’s t test or ANOVA followed by Tukey’s multiples comparisons test. Significance is expressed as *p<0.05, **p<0.01, ***p<0.001, ****p<0.0001.

## Supplementary figures

**Supplementary figure 1.**
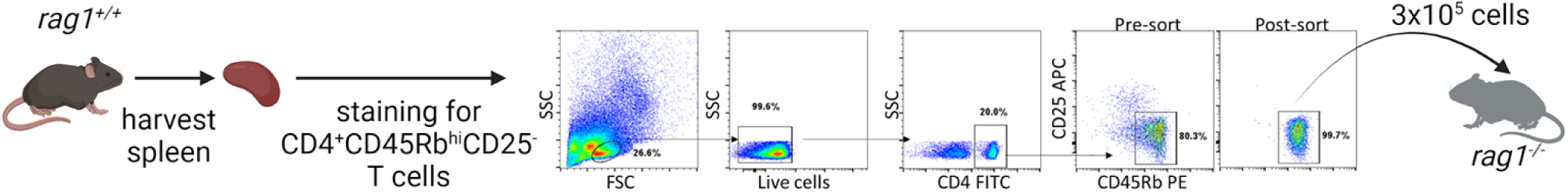
Gating strategy and the purity of naïve T cells. WT spleens were harvested and stained for CD4^+^CD25^-^CD45Rb^hi^ cells and were FACs sorted and a purity greater than 99% was confirmed before the transfer.

**Supplementary figure 2.**
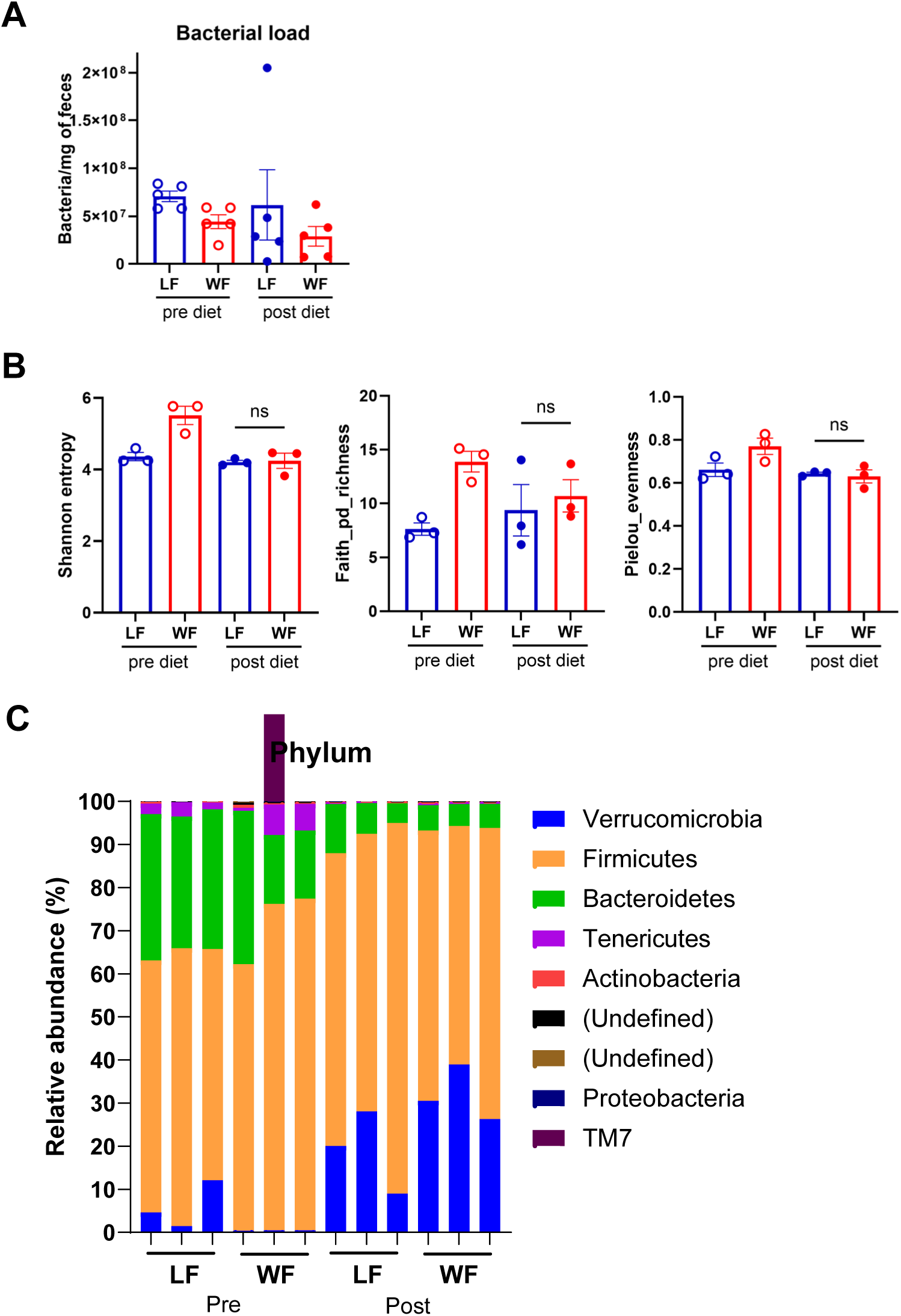
Fecal bacterial analysis of *Rag1^-/-^* mice on low-fiber or wheat fiber diets. *Rag1^-/-^* mice were fed either LF or Wheat for 2 weeks and their feces were analyzed. (A) Total bacterial load measured by qPCR with 16S rRNA primers. (B) Alpha diversity index analyzed by MiSeq 16S sequencing. (C) Taxonomic composition at the phylum level.

**Supplementary figure 3.**
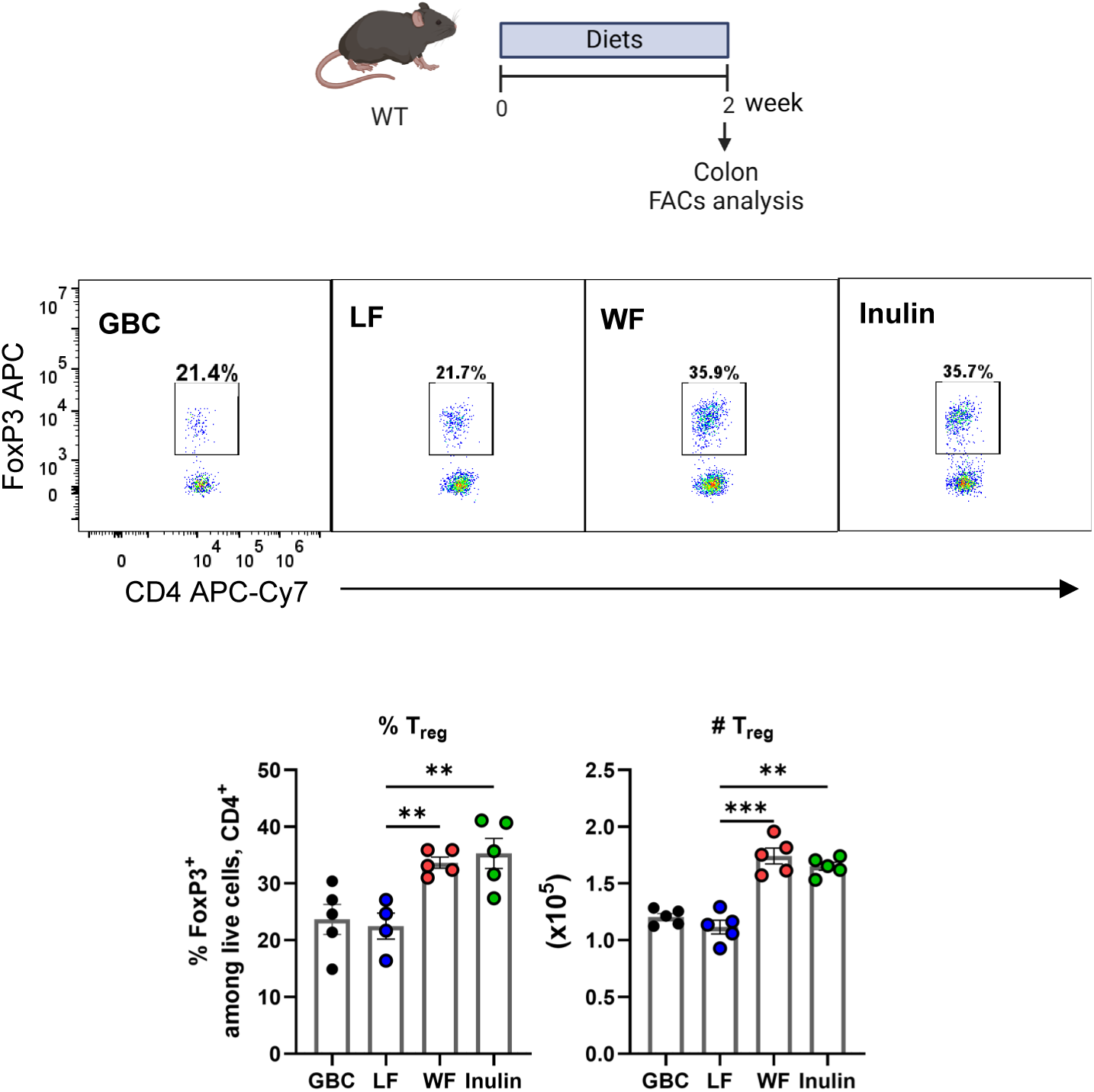
Wheat fiber’s induction of Tregs is comparable to that of fermentable fiber, inulin. WT conventional mice (n=5/group) were fed with LF, Wheat, or Inulin for 2 weeks and their colonic lamina propria Tregs were analyzed by flow cytometry.

## Notes

### Competing Interest Statement

The authors have declared no competing interest.

